# An efficient electroporation protocol for the genetic modification of mammalian cells

**DOI:** 10.1101/073387

**Authors:** Leonardo Chicaybam, Camila Barcelos, Barbara Peixoto, Mayra Carneiro, Cintia Gomez Limia, Patrícia Redondo, Carla Lira, Flávio Paraguassú-Braga, Zilton Vasconcelos, Luciana Barros, Martin Hernán Bonamino

## Abstract

Genetic modification of cell lines and primary cells is an expensive and cumbersome approach, often involving the use of viral vectors. Electroporation using square wave generating devices, like Lonza´s Nucleofector, is a widely used option, but the costs associated with the acquisition of electroporation kits and the transient transgene expression might hamper the utility of this methodology. In the present work we show that our in house developed buffers, termed Chicabuffers, can be efficiently used to electroporate cell lines and primary cells from murine and human origin. Using the Nucleofector II device, we electroporated 14 different cell lines and also primary cells, like mesenchymal stem cells and cord blood CD34+, providing optimized protocols for each of them. Moreover, when combined with Sleeping Beauty based transposon system, long-term transgene expression could be achieved in all types of cells tested. Transgene expression was stable and did not interfere with CD34+ differentiation to committed progenitors. We also show that these buffers can be used in CRISPR-mediated editing of *PDCD1* gene locus in 293T and human peripheral blood mononuclear cells. The optimized protocols reported in this study provide a suitable and cost-effective platform for the genetic modification of cells, facilitating the widespread adoption of this technology.

## Introduction

Cell lines are valuable tools for research development, constituting one of the pillars of experimental biology. Their unlimited proliferative capacity, high degree of homogeneity and relatively easy maintenance in culture allow the generation of large number of cells required for testing numerous candidate drugs (Barretina et al., 2012), ‐omics profiling (Blower et al., 2007; Griffin and Shockcor, 2004; Nishizuka et al., 2003) and signaling pathways studies (Park et al., 2010), to cite some examples. One of the areas that benefited the most with the use of cell lines was cancer research, with the derivation of several cell lines that can be used as models for different cancers. These cells are used to model disease in vitro and in vivo, providing information about oncogenesis related pathways and insights into therapeutic strategies (Gillet et al., 2013).Moreover, cell lines are central players in the biotechnology industry, being used in the production of biopharmaceuticals like antibodies, hormones, and bioactive proteins in general (Kuystermans and Al-Rubeai, 2015).

The use of cell lines in basic research is often associated with genetic modification protocols, which allow overexpression and/or silencing of desired genes in a controllable fashion. Recently, the development of gene editing tools like TALENs and CRISPRs provided a more precise control of gene insertion or deletion, extending the possible genomic manipulations (Kim and Kim, 2014). Methods to deliver foreign genetic material (DNA or RNA) usually rely in non-viral or viral vectors, with the former being preferred because of increased biosafety, easier production and faster translation. Electroporation is a non-viral method for gene transfer that is demonstrating encouraging results, being successfully used for the manufacture of antitumor lymphocytes (Ramanayake et al., 2015) and other applications (Kotnik et al., 2015), but the mechanism of DNA/RNA transfer is not fully understood (Satkauskas et al., 2012). Moreover, the use of electroporation is associated with extensive testing of electric parameters (pulse amplitude, volts) in order to optimize the protocol. Non-viral methods like liposomes and electroporation show varying efficiencies, with several cell lines and primary cells showing poor transfection rates and cell death (Wang et al., 2012; Yin et al., 2014). In the case of liposomes, the transfection of non-adherent cell lines is rather inefficient, showing good results only for some adherent cells (Behr, 2012; Jordan and Wurm, 2004).

Using a square wave pulse technology, Lonza´s Nucleofector electroporator was shown to be very efficient in several cell lines and primary human and murine cells, inducing high expression of the transgene and substantial viability24. The pre-loaded electroporation programs suited for each cell line simplify the experimental setup, and the use of proprietary additives improves the transfection efficiency. However, the frequent use of Nucleofector electroporation kits implies in important costs for research labs, especially those in middle to low income countries. In a previous work, our group developed “in house” electroporation buffers (termed “Chicabuffers”) that had comparable efficiency with Lonza´s buffers for the transfection of the human T cell line Jurkat and primary T lymphocytes from mouse and human origin(Chicaybam et al., 2013). Electroporation strategies using Chicabuffers were recently successfully applied to colon cancer cell lines (de Souza et al., 2013) and human mesenchymal stem cells (MSC; unpublished data). In the present work we extend the efficiency analysis of Chicabuffers and the description of optimal electroporation conditions in a panel of cell lines and primary cells that represent relevant models for cell biology studies and disease comprehension. We selected 14 cell lines of mouse and human origin and primary human cells (MSC, peripheral blood mononuclear cells -PBMCs -and cord blood CD34+ cells),showing that these buffers yield high transfection efficiencies and are a viable option for genetic modification using the Nucleofector IIb electroporator. For cells in which the levels of transgene expression was low, we developed SB-based transposon plasmids engineered to confer drug resistance, allowing fast and efficient drug based selection of cells representing fractions of the cell culture.

We selected cells lines representing models for hematopoietic neoplasias (HEL, K562, P815, Nalm-6 and Jurkat cell lines) and different solid tumor derived cell lines (A549, B16-F10, HeLa, MCF-7, MDA-MB-231). Some of the tested cells represent classical cellular models for ectopic gene expression (293T, NIH-3T3), cell signaling (Jurkat and 293T), growth factor dependence (BA/F-3) or simply relevant cells in terms of therapy and cell differentiation (MSCs, PBMCs and Cord Blood CD34+ cells). In addition, we show that the level of transfection achieved using Chicabuffers allows efficient genomic edition of the potentially clinical relevant PD1 locus in human cells such as 293T and peripheral blood mononuclear cells (PBMCs) using the recently described CRISPR/Cas9 system (Jinek et al., 2012).

## Materials and Methods

### Ethics approval

The use of PBMCs and CD34+ cells from healthy donors was approved by an IRB (Brazilian National Cancer Institute -INCA -Ethics Committee – protocol 153/13) and donors signed review board approved informed consents. MSCs were obtained from healthy donors submitted to surgery for hernia repair at the Clementino Fraga Filho University Hospital. The patients provided written informed consent and the study was approved by the Hospital Research Ethics Committee.

### Plasmids and Cloning

The pT3-GFP plasmid (Peng et al., 2009) was kindly provided by Dr. Richard Morgan (Surgery Branch -NCI). The pT2-GFP and SB100X (Mátés et al., 2009) constructs were kindly provided by Dr. Sang Wang Han (UNIFESP, Brazil). For the creation of pT3-Neo-EF1a-GFP plasmid, GFP was excised from pT3-GFP by digestion with AgeI/NotI and the neomycin resistance gene (NEO), which was synthesized by Genscript (Piscataway, NJ, USA), was inserted. The EF1a-GFP cassette was isolated from the plasmid pRRLsin.PPTs.EF1a.GFPpre (Bonamino et al., 2004) (provided by Dr. Didier Trono, EPFL, Switzerland) after digestion with ClaI/BstBI and inserted in pT3-NEO previously digested with ClaI. For CRISPR experiments, the plasmid encoding S. pyogenes Cas9 (WT) and a U6 promoter for guide RNA (gRNA) expression was acquired from Addgene (pX330; #42230). gRNA (5´CACCGGCCATCTCCCTGGCCCCCA 3´)for Programmed Cell Death 1 (*PDCD*-1) was designed by Optimized CRISPR Design tool (http://crispr.mit.edu/) and cloned in pX330 (Addgene) using BbsI restriction site. pRGS-CR (Kim et al., 2011) was provided by Dr Amilcar Tanuri (Federal University of Rio de Janeiro, Brazil), and *PDCD1* target sequence cloned in EcoRI / BamHI sites, between a red fluorescent protein (RFP) and a GFP, resulting in an out-of-frame GFP. The GFP expression can be restored by CRISPR-mediated non-homologous end joining (NHEJ) repair. All plasmids were isolated using Qiamp Maxi prep kit from Qiagen (Germany) and quantified using a Nanodrop spectrophotometer. The new constructs described in this report are available at Addgene.

### Cell lines and primary cells

The origin and cell culture conditions for each cell line are described in Table S1. The use of PBMCs from healthy donors was approved by an IRB (Brazilian National Cancer Institute -INCA -Ethics Committee) and donors signed review board approved informed consents. Within 24h after blood collection, leukocytes were harvested by filtration and washed with Phosphate Buffered Saline (PBS). A density gradient centrifugation using Ficoll-Hypaque^®^-1077 was performed. Cells were centrifuged for 20min at 890g (slow acceleration/deceleration off), washed three times with PBS and used for nucleofection. For CD34+ cells separation, mononuclear cells (MNCs) were isolated from umbilical cord blood after Ficoll density gradient using the same protocolabove described. CD34+ cells were isolated from MNCs using CD34 MicroBead Kit (Miltenyi Biotech) following the manufacturer’s instructions. The utilization of CD34+ cells was also approved by INCA’s Ethics Committee.

MSCs were isolated from abdominal subcutaneous adipose tissue fragments obtained from healthy donors submitted to surgery for hernia repair at the Clementino Fraga Filho University Hospital. The patients provided written informed consent and the study was approved by the Hospital Research Ethics Committee. Fragments were cut into small pieces and incubated with 1 mg/ml collagenase type II (Sigma-Aldrich, MO, USA)under permanent shaking at 37°C for 30 minutes. The cell suspension was centrifuged at400 g, room temperature, for 10 minutes and the pellet was resuspended on PBS, followed by filtration with 100 μm mesh strainers. Cells were plated to expand MSCs at3×104 cells/cm2 density with low-glucose Dulbecco’s modified Eagle’s medium (DMEM Low-glucose, Gibco, CA, USA) supplemented with 10% fetal bovine serum (Gibco, CA, USA) and 100 U/ml penicillin and 100 μg/ml streptomycin (Sigma-Aldrich, MO, USA).Cells were electroporated at passage 3.

### Electroporation

Generic cuvettes were used for all the electroporations (Mirus Biotech^®^, Madison, WI, USA cat.: MIR 50121). Cells were resuspended in 100ul of the desired buffer and 4ug of the reporter plasmid (pT2-GFP transposon) were added. For long term experiments, 1ug of SB100X was added. The seven different buffers tested in this work are described in Table S2. Cells were transferred to a sterile 0.2cm cuvette and electroporated using the reported program (Table 1) of Lonza^®^ Nucleofector^®^ II electroporation system. After transfection, cells were gently resuspended in 1mL of pre-warmed RPMI medium supplemented only with 2mM L-Glutamine and 20% FCS. All cells were seeded in 12-well plates and grown at 37^o^C and 5% CO2. The medium was replaced by complete RPMI medium the following day and cells were maintained as described previously.

**Table 1:** Summarized electroporation conditions for each cell line (based in figure 2; d1 after electroporation).

| Cell Line | Cell Type | Program | Recommended Buffer | Viability (Chicabuffer; ±SD) | Viability (Lonza) | GFP expression (Chicabuffer) | GFP expression (Lonza) |
| --- | --- | --- | --- | --- | --- | --- | --- |
| <u>Non-adherent</u> |  |  |  |  |  |  |  |
| BA/F3 | Mouse pro B cell | X-001 | 2M | 91,8±2,7 % | 79,00 % | 55,05±14,2 % | 88,00% |
| HEL | Erythroleukemia; erythroblast cell | X-005 | 1S | 70,4±17,1 | 39-66% | 79±6,2% | 94,00% |
| Jurkat | Acute T cell leukemia, T lymphocyte; lymphoblastoid cells | X-001 | 1SM | 75,7±6,8 % | 90,00 % | 69±11,6 | 88,00% |
| K562 | Human chronic myelogenous leukemia; lymphoblastoid cells | T016 | 1M | 70,7±13,8 % | 88,00 % | 64,1±8% | 80-90% |
| Nalm-6 | Human B cell precursor leukemia | C-005 | 3P | 74,2±11,8 % | 87,00 % | 40,6±14,7 % | 64,00% |
| P815 | Mouse mastocytoma; mast cells | C-005 | 3P | 70,2±29,9 % | 92,00 % | 60,5±16,6 % | 62,00% |
| <u>Adherent</u> |  |  |  |  |  |  |  |
| A549 | Human lung carcinoma; epithelial cells | X-001 | 3P | 59,4±27,3 % | 81,00 % | 63,5±11,4 % | 72,00% |
| 293T | Human embryonal kidney; adherent fibroblastoid cells | A-023 | 1SM | 79,9±24,7 % | 90,00 % | 38,6±30,2 % | 90,00% |
| B16F10 | Mouse skin melanoma | P-020 | 2S | 39,9±17 % | 91,00 % | 49,3±15,7 % | 84,00% |
| HeLa | Human cervix carcinoma; epitheloid cells in monolayers | I-013 | 1M | 45,1±16,5 % | 85-90% | 66,4±8,3 % | 70,00% |
| MCF7 | Human breast adenocarcinoma; epithelial cells | P-020 | 1M | 68,4±10,9 % | 60,00 % | 57±23,3 % | 77,00% |
| MDA-MB-231 | Human breast adenocarcinoma; epithelial cells | X-013 | 1SM | 85,6±15 % | 77,00 % | 48,5±17,5 % | 79,00% |
| human MSCs | Human mesenchymal stem cells | U-023 | 2S | 58,5±6,8 % | 48,00 % | 35±16,6 % | 80,00% |
| NIH3T3 | NIH Swiss mouse embryo; adherent fibroblastoid cells | U-030 | 1SM | 49,4±30,4 % | 87,00 % | 52,5±19,5 % | 84,00% |

### Electroporation Score determination

For non-adherent cell lines, viability determination was based on trypan blue exclusion and/or determination of the % of cells displaying viable cell FSC vs SSC parameters by flow cytometry analysis on cells negative after 7AAD staining. For adherent cells, viability determination was calculated based on the % of the OD obtained in Crystal Violet staining assays at d+1 or d+3. Calculation was based on the formula %= 100x [OD for control (non electroporated) cell line/(OD for control (non electroporated)cell line + OD for electroporated cell line). The "electroporation score" was calculated based on cell viability (after normalization against the viability of non-transfected cells)and transgene expression on d+1, and the score set to the formula “Viability(%)*Expression (%)/F”. A division factor (F=50 for adherent cell lines and F=100 for non-adherent cell lines) was used in the score formula to fit the results in the graph scale.

### Crystal Violet staining

To assess viability of adherent cell lines, cells were plated in triplicate in 96-well microtiter plates immediately after electroporation. Cell viability was evaluated after 24 hours and cell expansion was analyzed at day+1 by crystal violet. The crystal violet incorporation assay was performed by fixing the cells with ethanol for 10 min, followed by staining them with 0.05% crystal violet in 20% ethanol for 10 min and solubilization with methanol as reported (Faget et al., 2012). The plate was read on a spectrophotometer at 595 nm (SpectraMax 190, Molecular Devices, Sunnyvale, CA).

### In vivo B16-F10 tumor model

B16F10 cells were electroporated with 4μg of pT3-NEO-EF1a-GFP and 1μg of SB100x in buffer 1S, program P-020 of Lonza Nucleofactor II. As negative controls, we electroporated cells only with pT3-NEO-EF1a-GFP. Each condition was plated in a 6-well plate. After reaching 80% confluence, G418 (Life Technologies) antibiotic was added at 2,000μg/mL. The medium was changed every three days and the antibiotic added. After selection with antibiotic or not, we injected 5x10^5^ cells in the left flank of C57BL/6 mice. After 15 days, we excised the tumor and plated the cells in 25cm^2^ culture flasks. After 24 hours, the culture medium was changed to eliminate non-adherent cells. After 3 days, the cells were recovered and analyzed by flow cytometry for GFP expression.

### CD34+ differentiation assay

Electroporated CD34+ cells were assayed in two different concentrations, 5x102 and 2x103 cells/well. The cells were concentrated in 300μL and then added in 1.1x concentrated 3 mL Methocult™ H4034 (Stem Cell Technologies Inc., Vancouver, Canada) then seeded 2 wells of a six-well plates, 1.1mL/well. Cells were cultivated for three weeks at 37 °C in a humidified atmosphere supplemented 5% CO2 in incubator 300/3000 Series (Revco, Ohio, EUA). The colonies were identified and quantified using STEMvision™ (Stem Cell Technologies Inc.) for the burst-forming units-erythroid(BFU-E), colony-forming units-erythroid (CFU-E), colony-forming units-granulocyte or macrophage or granulocyte-macrophage (CFU-G/M/GM) and colony-forming units-granulocyte/erythroid/megakaryocyte/macrophage (CFU-GEMM).

### Flow cytometry

FACSCalibur^®^ (BD Bioscience) was used to perform morphologic evaluation of viability (FSC vs SSC) and GFP expression analysis. Cells were harvested the following days after transfection and resuspended in PBS at a concentration of 105 cells/500uL. 7AAD staining (eBioscience cat. 00-6693) was performed immediately before FACS acquisition following manufacturer instructions. Data were analyzed using the FlowJo software (Tree Star). The hematopoietic progenitor CD34+ cells were evaluated for purity by staining with anti-CD34-PE (clone 581, BD Biosciences).

### CRISPR-mediated gene editing

HEK293FT and PBMCs were electroporated with pX330-PDCD-1 (10μg) and pRGS-CR-target (5μg). Gene editions were evaluated by GFP+/RFP+ ratio after 24 hours by flow cytometry. To characterize indels at *PDCD*1 locus, genomic DNA of gene edited cells was isolated by phenol-chloroform. Amplification of the target region was performed by PCR using the forward 5’-CCCCAGCAGAGACTTCTCAA and the reverse 5’-AGGACCGGCTCAGCTCAC primers. The PCR fragment was ligated inpCR2.1 vector (TA Cloning^®^ Kit, Life Technologies), transformed in DH5α cells and single bacteria colonies has the plasmid DNA extracted and sequenced using the primers described above.

### Short RNA and plasmid co-electroporation

After Ficoll gradient purification, PBMCs (107 cells) were electroporated with pRGS-CR-target (10μg) and 10–50pmol of FITC labeled RNA (Invitrogen) in Chicabuffer 3P and U-014 Nucleofector IIb program. Cells were left resting in RPMI+10%FCS for 24h at 37ºC and 5% CO2 and then evaluated by flow cytometry using ACCURI C6 (BD Bioscience).

### Statistical analysis

Data from electroporation experiments were analyzed by one-way ANOVA followed by Tukey´s multiple comparison test using GraphPad Prism 6 software.

## Results

With the objective of determining the best-suited buffer for the electroporation ofeach cell line, cells were electroporated with seven different buffers and the viability and GFP expression were analyzed. Representative flow cytometry plots are depicted in figure 1, showing 7AAD staining and GFP signal (gated in 7AAD negative cells) for a high electroporation score cell line (HEL) and FSC/SSC and GFP signal for a low score cell line (NIH3T3). 7AAD staining was performed only in the non-adherent cells since they represent a mixture of viable and non-viable cells at day 1 post electroporation. Adherent cells were allowed to adhere overnight after electroporation and non-adherent/dead cells were discarded before FACS analysis. As showed in figure 2, the majority of cell lines showed high electroporation scores independent of the buffer, with exception of P815, which showed an overall low efficiency but demonstrated best performance with buffer 3P. Suspension cell lines showed the best results regarding GFP expression, in which values above 60% were recurrently obtained. One exception is Nalm-6, with a maximum of 40% of GFP-positive cells obtained using buffer 3P. Adherent cell lines showed GFP values slightly lower (30–65%), with Hela showing the best result with 66,4±8,3% of GFP expression using buffer 3P. Importantly, after 24h of electroporation the cells showed a good viability (Fig. 2), allowing expansion and recovery from the nucleofection. Viability and GFP expression were followed for 10 days (suspension cell lines) or 7 days (adherent cell lines), with some cells retaining high levels of GFP (K562, HEL, B16) and others showing low expression of the marker after the expansion (NIH3T3, Jurkat, P815) (Figure S1-14). These results probably reflect the observed differences in nucleofection efficiency and proliferation rates among the studied cells. The electroporation protocol for each cell line is summarized in Table 1.

**Figure 1:**
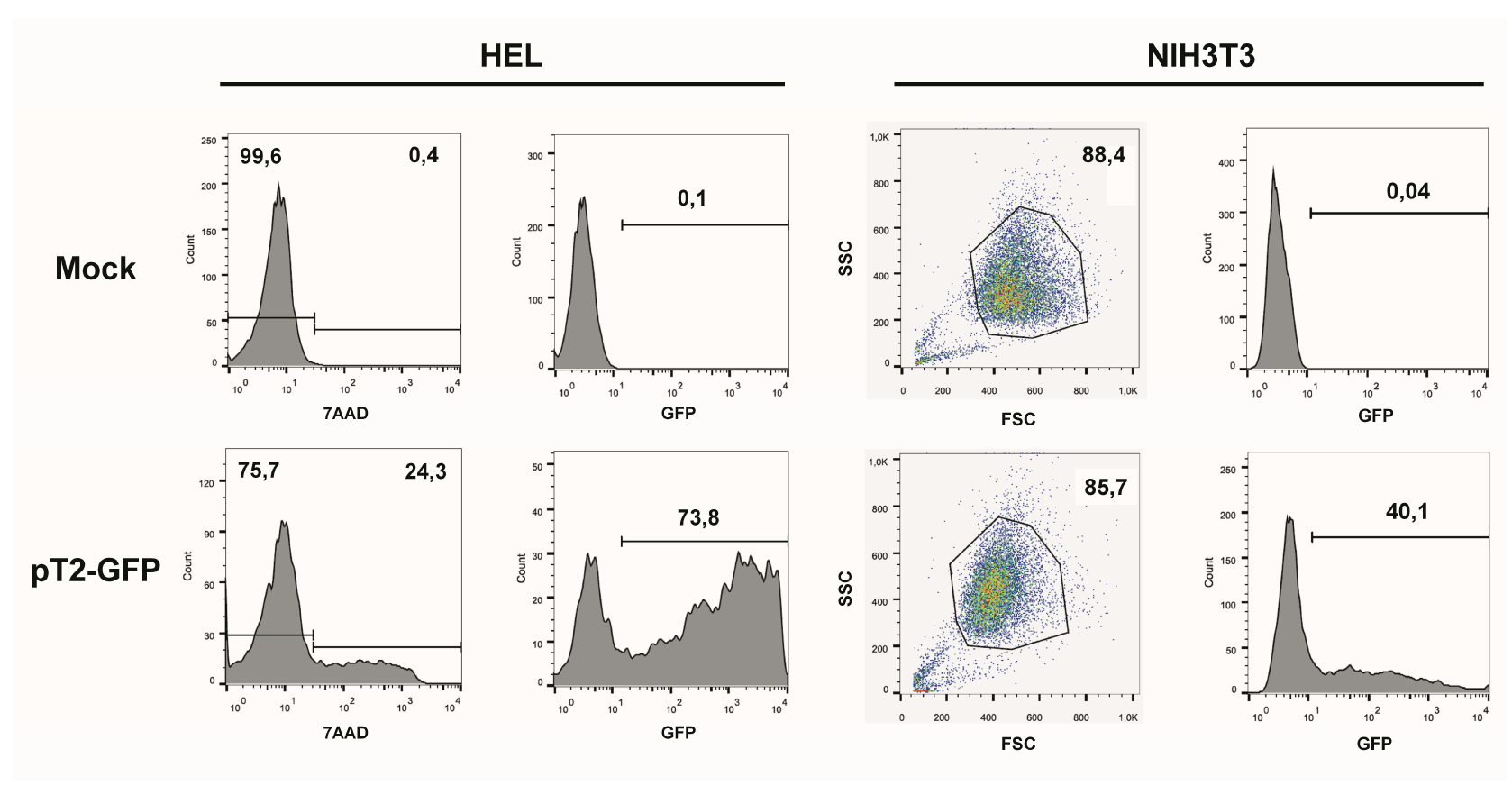
GFP expression after electroporation of representative cell lines. Representative plots of a high score cell line (HEL) and a low score cell line (NIH3T3). HEL was electroporated using buffer 2S and program X-005 and NIH3T3 using buffer 1SM and program U-030. For HEL, on day one after nucleofection cells were stained with 7AAD (left column of graphs) and GFP expression was analyzed on 7AAD negative population (right column). For NIH3T3, viable cells were gated based in FSC/SSC and GFP was analyzed. Numbers depict the percentages of cells in each gate.

**Figure 2:**
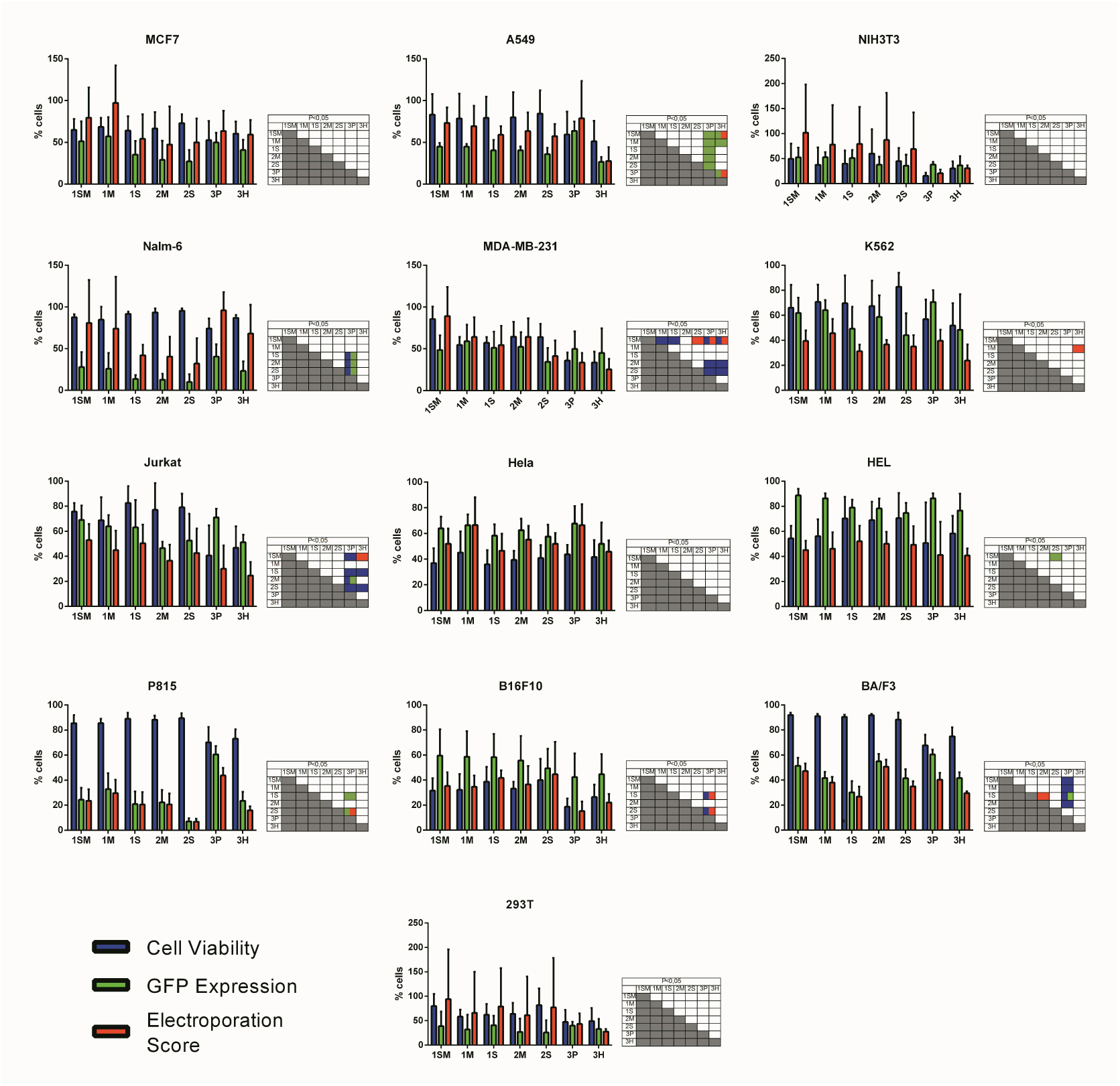
Electroporation score for cell lines. Cell lines were electroporated with pT2-GFP (4ug) using each one of the seven buffers and the recommended program. Viability (blue bar), GFP expression (green bar) and electroporation score (red bar) were assessed one day after nucleofection (d+1). Viability data were normalized with viability from non-transfected cells. Data are shown as mean ± SD from three experiments performed in duplicate and were further analyzed using one-way ANOVA with Tukey´s multiple comparisons test. Significant differences (p<0,05) are depicted in the table next to each graph, with each color denoting one parameter.

Stable gene expression is often required in the experimental setting, allowing the generation of subclones with overexpression or silencing of a gene of interest. The emergence of nonviral vectors that allow the integration of transgenes, like the Sleeping Beauty (SB) transposon system, simplified the genetic modification of cells, requiring only the delivery of two plasmids to achieve stable expression (one encoding the transgene flanked by ITRs – inverted terminal repeats – and one encoding the transposase). In order to evaluate if Chicabuffers could be used with this system, 1ug ofSB100x (encoding a hyperactive version of the SB transposase) was electroporated with4ug of pT2-GFP and GFP expression was followed for 30 days. As showed in figure 3, the addition of the SB100x induced a higher percentage of GFP-positive cells after 30 days of culture when compared with control cells, strongly suggesting that integration ofthe transgene has occurred. This effect was more pronounced in B16F10, HeLa and MCF7 cell lines, with approximately 20% of GFP-positive cells at day 30. The other celllines showed only a modest increase in GFP-positive cells at day 30, ranging from 2% (BA/F-3) to 12% (K562). The long-term levels of GFP expression did not correlate with GFP expression at early days after nucleofection, suggesting that the cell lines have different intrinsic susceptibilities to SB-induced transgene integration.

**Figure 3:**
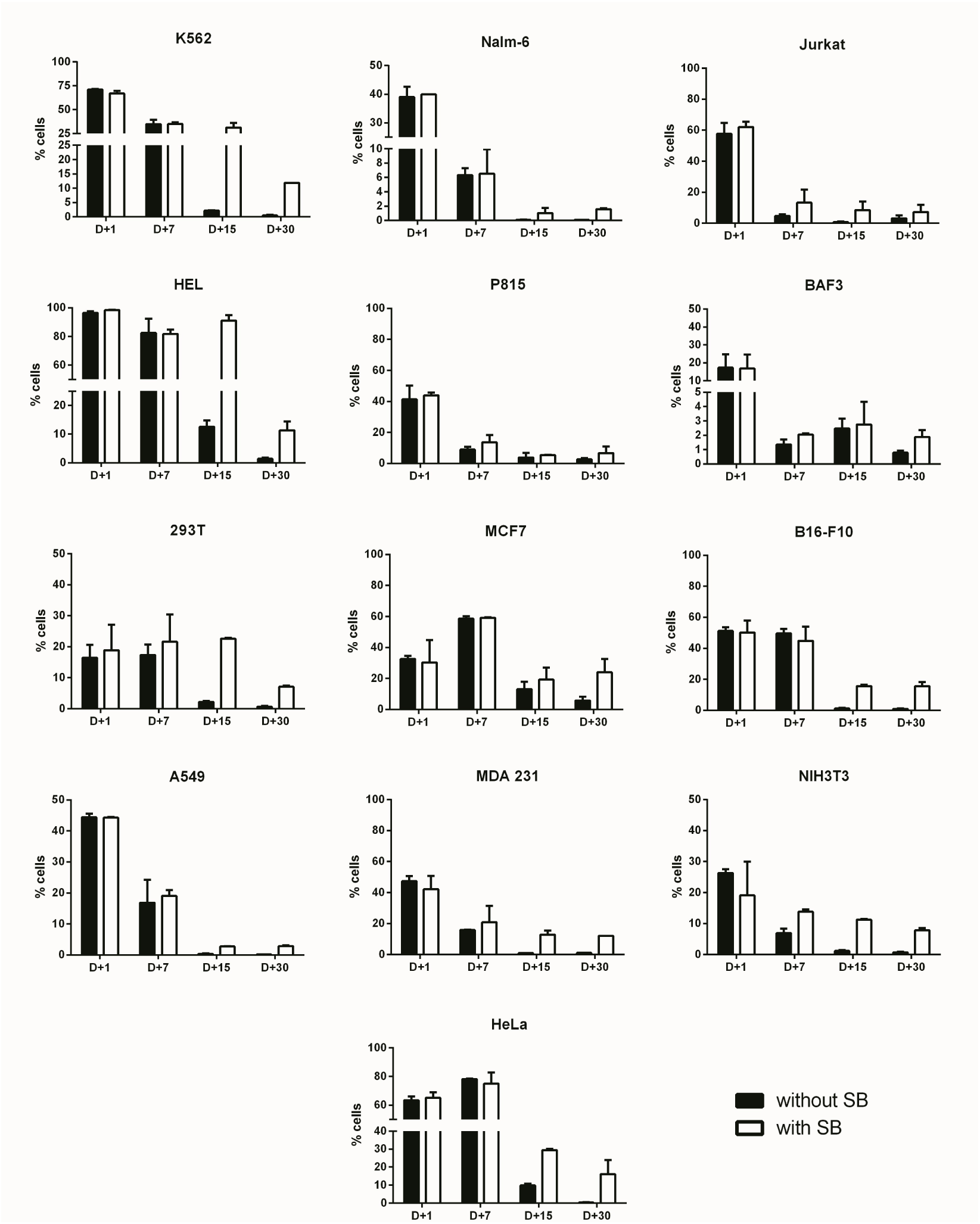
Long term transgene expression in electroporated cell lines using SB system. Cell lines were electroporated with pT2-GFP (4ug) using the combination of buffer and program indicated on table 1, with (white bar) or without (black bar) the addition of SB100x transposase (1ug). GFP expression was analyzed until d+30 for each cell line. Data are shown as mean ± SD from one single experiment performed in duplicate.

For fast and easy enrichment of GFP-positive cells we constructed a bidirectional vector encoding GFP and G418 resistance in the backbone of pT3 transposon, named pT3-Neo-EF1a-GFP. Indeed, the expression level obtained after nucleofection was sufficient to select G418-resistant clones after electroporation with this plasmid, as shown for NIH3T3 (Fig. 4A) and B16F10 (Fig. 4B) cell lines. After G418 selection and withdrawal, GFP expression remained stable in NIH3T3 cells for 15 days (Figure S15).Furthermore, when the modified B16F10 cells were injected in vivo and allowed to form subcutaneous tumors, the cells extracted from the tumor at d+14 post inoculation (dpi)still expressed high levels of GFP, indicating that the transgenic cassette is integrated inthe genome and has stable expression, with no signs of in vivo silencing of the transgene (Figs. 4C and Figure S16).

**Figure 4:**
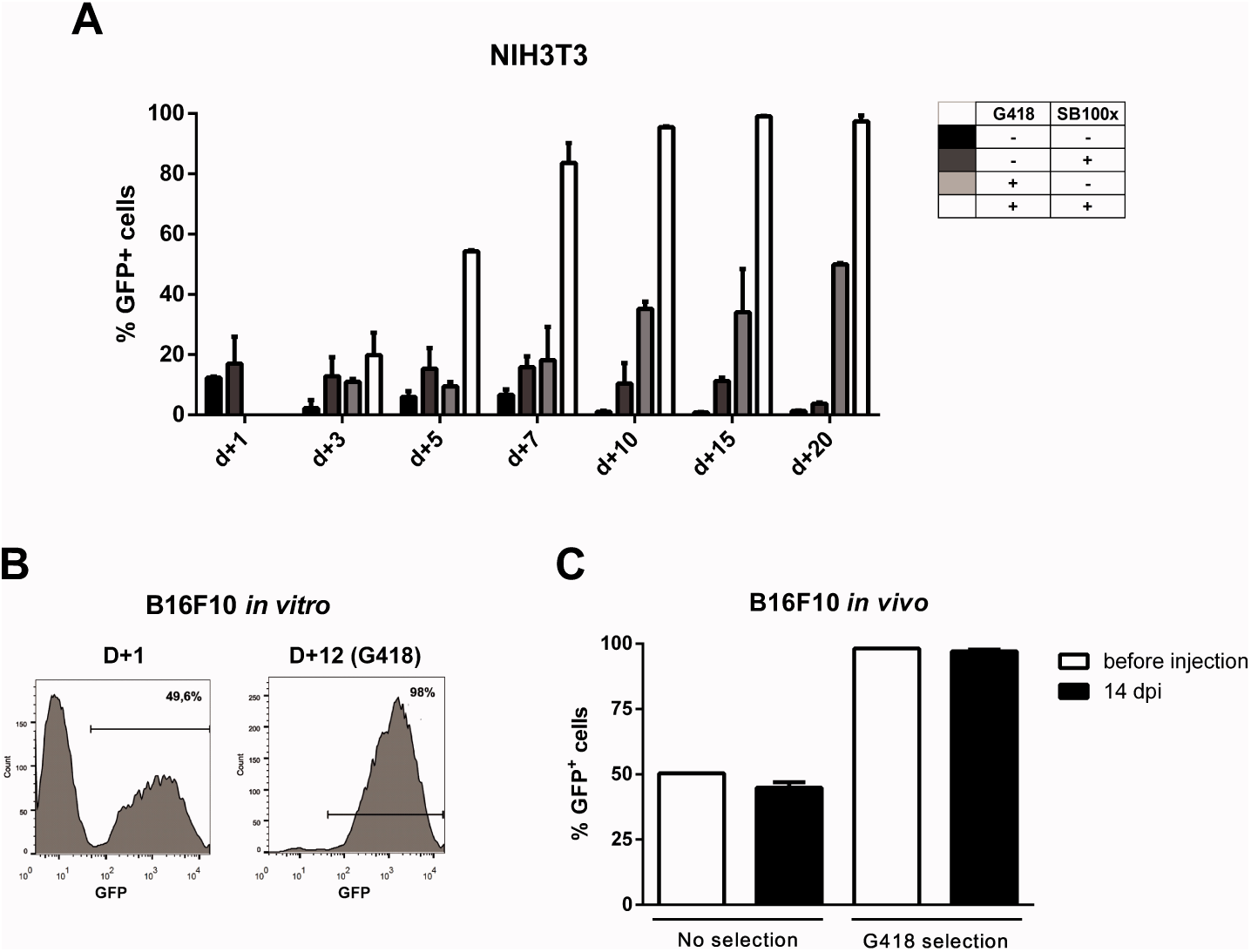
Transgene expression can be enriched by using G418 and is retained after *in vivo* growth. Using the programs and buffers indicated on Table 1, NIH3T3 (**A**) and B16F10 (**B**) cell lines were electroporated. G418 was added two days after nucleofection and GFP expression was accompanied until d+20 (NIH3T3) or d+12 (B16F10). (**C**) 5x10^5^ B16F10 cells submitted or not to selection with G418 were injected in the left flank ofC57Bl/6 mice. Tumors were extracted 14 days post injection (dpi), cells were passed in vitro for one week and GFP expression analyzed by FACS.

Variations of the pT3-Neo-EF1a-GFP construct were developed, such as the pT3-Neo plasmid, which confers resistance to G418 antibiotic and has restriction sites that allow cloning of a second expression cassette. This plasmid was validated in G418 resistance assays using B16F10 cells (data not shown). The map for this plasmid is shown in Figure S18.

The use of primary cells derived from patients or healthy donors provides a more accurate model for in vitro and in vivo experiments, and these cells can also be used in cell therapy approaches to treat a large number of diseases. However, these applications often depend on genetic modification, which is usually hard to perform in these cells. To evaluate the performance of Chicabuffers in the gene transfer to these cells, we isolated adipose tissue derived MSCs and cord blood purified CD34+ hematopoietic stem cells and electroporated the cells with the plasmids pT2-GFP and SB100x. As shown in figure 5A, the best electroporation score for MSC was obtained using buffer 2S, with 57% of viable cells and 39% of GFP expression. When using SB100X, long-term expression ofGFP using this buffer was seen in 12% of cells (Fig. 5B). For CD34+ cells, around 57% were GFP-positive one day after electroporation using buffer 1SM and program U-008 (Fig. 6A). These cells were plated in methylcellulose-based medium, allowing long-term assessment of GFP expression and differentiation potential. After three weeks, GFP+ CD34+ cells were able to differentiate to erythroid, granulocytic and myeloid lineages (Fig. 6B), showing that the insertion of the transgene did not affect the stemness of the cells and that differentiated cells display high GFP expression (Figure S17).

**Figure 5:**
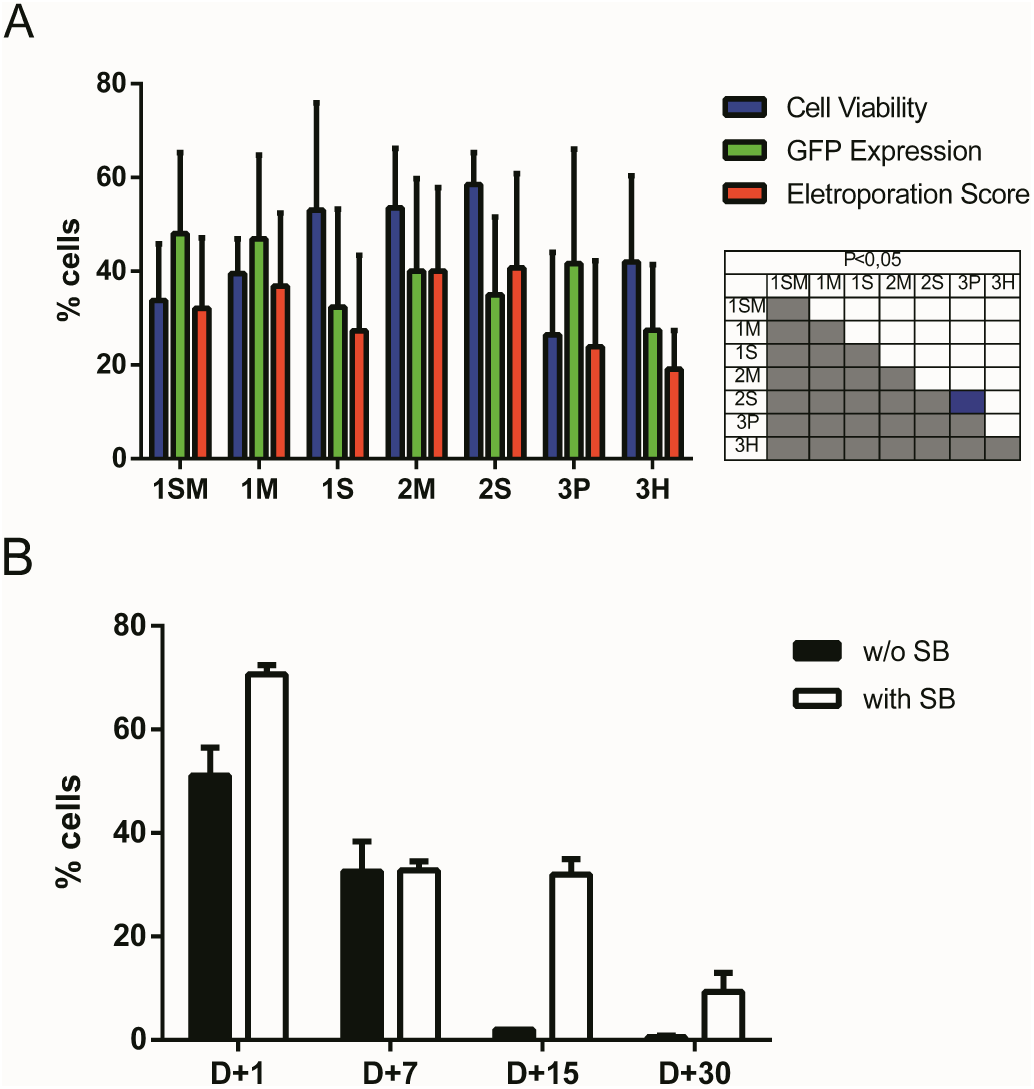
SB based GFP gene transfer to adipose tissue derived human MSCs. (**A**)MSCs were electroporated with each one of the seven buffers and the recommended program. Viability (blue bar), GFP expression (green bar) and electroporation score (red bar) were assessed one day after nucleofection (d+1). (**B**) Long term GFP expression was evaluated until d+30 post nucleofection with (white bar) or without (black bar) the addition of SB100x transposase (1ug per cuvette). Data are shown as mean ± SD from three experiments performed in duplicate and were further analyzed using one-way ANOVA with Tukey´s multiple comparisons test. Significant differences (p<0,05) are depicted in the table next to each graph, with each color denoting one parameter.

**Figure 6:**
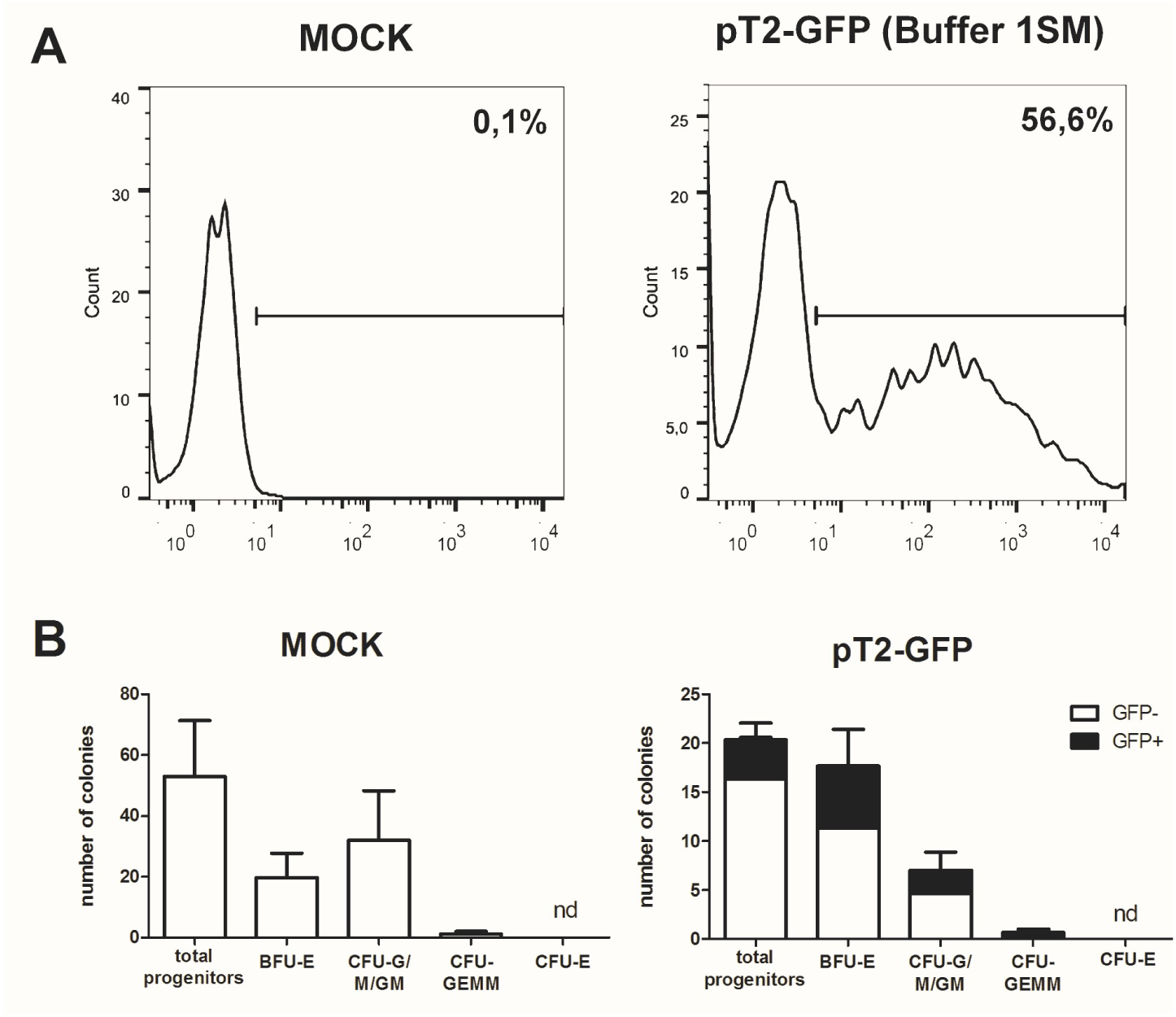
SB based GFP gene transfer to human cord blood CD34+ cells. (**A**) GFP expression in CD34+ cells electroporated with plasmids pT2-GFP (4ug) and SB100x (1ug) using program U-008 and buffer 1SM. GFP expression was evaluated by FACS atd+1 post nucleofection. (**B**) Electroporated cells (2x10^3^ per well) were plated in methylcellulose media and allowed to differentiate for 3 weeks. Colonies were quantified for mock (left) and GFP electroporated (right) cells. GFP positive colonies (black bars)were determined within total colonies identified. CFU-E: colony forming unit – erythroid;BFU-E: burst forming unit – erythroid; CFU-G/M/GM: colony forming unit – granulocyte/monocyte/; CFU-GEMM: colony forming unit – granulocyte-erythrocytemonocyte-megakaryocyte. Data are shown as mean ± SD from two experiments.

The recent description of the CRISPR/Cas9 system as an efficient tool to edit the genome of cells has clear implications for basic cell biology studies and gene therapy protocols (Doudna and Charpentier, 2014). To achieve efficient gene editing of target cells, Cas9 nuclease and the guide RNA (gRNA) must be expressed in the cell, ideally in a transient fashion. To evaluate the efficiency of Chicabuffers in promoting Cas9 mediated genome editing, we designed a gRNA targeting exon 2 of *PDCD*1 gene, which encodes the inhibitory receptor PD-1, a relevant potential target for cancer cell based immunotherapy (Chicaybam and Bonamino, 2014; Hamid et al., 2013). For the validation of gRNA, we used plasmid pRGS-CR-PDCD1, which has the *PDCD1* target sequence cloned between a red fluorescent protein (RFP) and a GFP, resulting in an out-of-frame GFP. In this system, GFP expression can be restored by CRISPR-mediated non-homologous end joining (NHEJ) repair (Kim et al., 2011), leading to restoration of the reading frame in nearly 1/3 of the editions. Co-electroporation of 293T cells with the report construct and the plasmid carrying CRISPR/Cas9/gRNA, but not CRISPR/Cas9 lacking the gRNA sequence, resulted in GFP expression in approximately 7% of the RFP+ cells (3% out of 42%), indicating that sequence-specific DNA editing was achieved (Fig. 7A). A similar approach was performed in PBMCs and following electroporation, indels were verified by amplification of *PDCD1* locus of the edited cells, which was subsequently cloned in pCR2.1 vector and analyzed by Sanger sequencing, evidencing cells containing indels of varying lengths in the *PDCD1* locus (Fig. 7B). The results of gene editing experiments in 293T and PBMCs are summarized in figure 7C. The characterization of indels in PBMCs and 293T cells indicate that the use of our optimized electroporation protocol allowed efficient editing of *PDCD*1 locus in the tested samples. All the indels led to disruptions of the reading frame of the PD1 sequence (data notshown).

**Figure 7:**
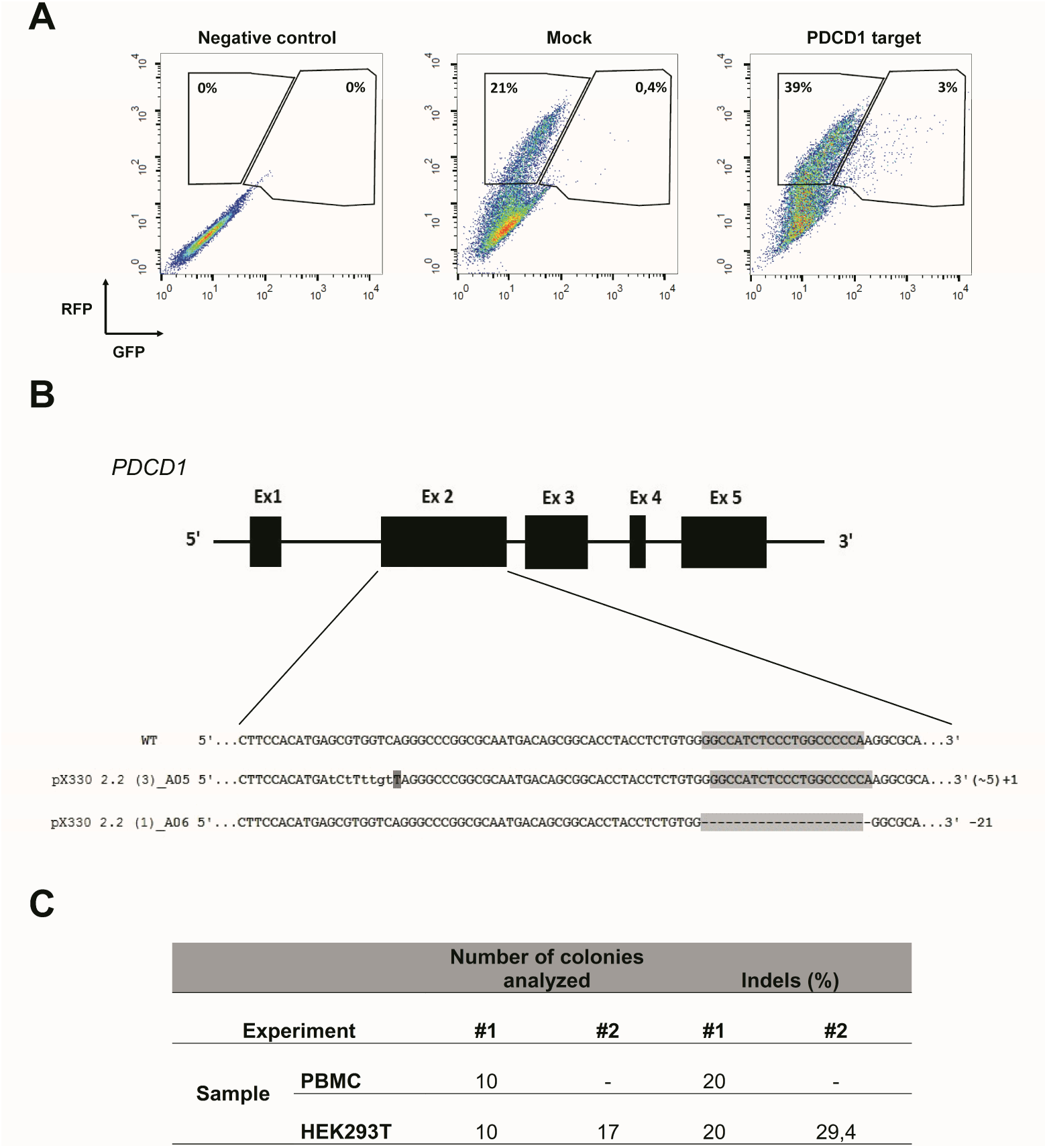
Electroporation of CRISPR/Cas9 cassettes promotes gene editing ofPBMCs and 293T cells. (**A**) 293T cells were electroporated (buffer 3P, program A-023)without plasmid (negative control), with pRGS-CR plasmid (without *PDCD1* target sequence; mock) or with pRGS-CR-PDCD1. GFP and RFP expression was analyzed 24 hours later. Numbers depict the percentage of cells inside each gate. (**B**) Representative image showing indels obtained in *PDCD1* gene after electroporation of PBMCs with plasmid px330 (Cas9/gRNA). The indels are represented by lower case characters;numbers inside parenthesis depict substitutions (~) and numbers outside parenthesis depict additions (+) or deletions (-). Exons are not draw into scale. (**C**) Summarized results obtained for 293T cells and PBMCs, showing the number of colonies sequenced and the percentage of indels detected. Two experiments were done for each cell.

Multiple target editing is possible using CRISPR systems. Since multiple loci editing require multiple gRNA, we evaluated the possibility of co-electroporating PBMCs with a reporter plasmid and FITC labeled short RNAs. This setting could be used to co-electroporate a plasmid encoding a reporter gene (or Cas9 nuclease) and multiple short RNAs (such as gRNAs for editing several loci). Using the buffer 3P we were able to achieve high viability (Figure S19a) and up to 60,7% of cells expressing the short RNA when 50pmol of the RNA were used (Figure S19b). Concentrations above 75pmol of short RNA resulted in increased cell death and were not further used (data not show).From electroporated cells under the same condition, up to 14,8% co-expressed the reporter plasmid (encoding RFP) and the labeled short RNA (Figure S19c). This setting clearly allows efficient co-electroporation of plasmid DNA and short RNA, opening the possibility of combining siRNA and transgene expression or even multiple gRNAs and Cas9 expressing plasmids for gene editing.

## Discussion

Genetic modification of cells is a cumbersome and expensive process, often involving the use of viral vectors to achieve high efficiency transgene expression. The use of electroporation for the genetic modification of cells is being adopted by many laboratories as it represents a fast and cheap option for transfer of plasmids and RNA. Moreover, this technique is also very efficient, inducing transgene expression levels comparable to viral vectors in some cells (Bilal et al., 2015). Equipments capable of generating square-wave voltage pulses, like Lonza Nucleofector, are among the most efficient for mammalian cell electroporation (Mir, 2014). However, costs associated with the acquisition of nucleofection kits, especially if used in a routine basis, might hamper the use of this technology in some laboratories or impair large-scale experiments.

In a previous work, our group described seven in house buffers and tested the electroporation efficiency of Jurkat cells and primary lymphocytes using Nucleofector (Chicaybam et al., 2013). The selected buffers induced high transgene expression and low toxicity, comparable to results obtained when Lonza´s kit were used. In this context, the present work comprises a practical guide for the electroporation of 14 cell lines and primary MSCs and HSCs, determining the best buffer (among seven options) to be used with Lonza Nucleofector II, a widely disseminated electroporation device. The electroporation score calculated for every cell line is a general guide for electroporation efficiency comparison, and the buffer choice can be adapted to the need of the planned experiment (higher GFP expression or cell viability). Chicabuffers showed to work for all the cells tested with most of the samples showing interchangeable results among the different buffers and only few exceptions where one of the buffers performed poorly. This results place Chicabuffers as a valuable tool for cheap and fast gene modification of basically every cell tested. Although we focused in Lonza´s device, it is likely that a similar approach using these buffers in conjunction with electroporators that allow modification of electroporation conditions could achieve even better results by fine tuning parameters like pulse amplitude, voltage and wave forms (Yarmush et al., 2014). Lonza`s buffers were already described to have good results when tested with alternative nucleofector IIb programs (Gresch et al., 2004), suggesting that there is still room for optimization of electroporation conditions, reinforcing the potential of testing Chicabuffers under different experimental settings.

Short-term viability and expression of GFP was very efficient for the majority of cell lines, and Chicabuffers performed equally well when compared to the results reported by Lonza, especially for non-adherent cell lines (Table 1 and Fig. 2). Furthermore, our results are comparable to those reported in the literature for cell lines like K562 (Gresch et al., 2004) primary MSCs (Aluigi et al., 2006), although direct comparison of the results must be taken carefully because different plasmids were used. By combining this strategy with the Sleeping Beauty transposon system, the provided optimized protocols allowed long-term expression of transgenes in all the cells tested (Fig. 3). In the case of viral vectors, especially retroviral and lentiviral vectors, there is a wide availability of constructs carrying selectable markers, fluorescent reporters, promoters for different finalities and cassette configurations, increasing the options of possible cellular manipulations (Szulc et al., 2006; Vargas et al., 2012; Weber et al., 2008). This is in sharp contrast to the Sleeping Beauty system, which has a limited offer of transfer plasmids available. The new vectors developed and validated in the present report can improve flexibility and increase the applicability of this system, promoting accessible and efficient transgene integration into different cell types. These plasmids showed high and stable levels of transgene expression, and the addition of antibiotic resistance allowed the selection of GFP-expressing clones in vitro. Long-term expression of the transgene can be potentially increased by the use of SB100x RNA, decreasing the toxicity of the electroporation process as reported (Peng et al., 2009), or by carefully titrating the transposase plasmid mass to avoid overproduction inhibition (Grabundzija et al., 2010).These vectors and others recently reported in the literature (Kowarz et al., 2015), in conjunction with Chicabuffers, could be potentially used in diverse experimental gene therapy approaches, such as T cell immunotherapy (Singh et al., 2015), MSC (Martin et al., 2014) and stem cell gene therapy protocols (Aiuti et al., 2013), further facilitating the application of these technologies in basic, translational and clinical studies.

Our results show the feasibility of this approach, enabling a stable transgene expression in CD34+ cells from cord blood samples, keeping GFP expression throughout hematopoietic differentiation. It would be interesting to test this strategy in stem cell differentiation models other than the hematopoietic system such as the central nervous system (Sartore et al., 2011), including models of in vivo differentiation. In addition, cells with clear therapeutic potential, such as T lymphocytes (Chicaybam et al., 2013) and MSCs (this report) could be stably modified using a combination of Chicabuffer, SB and electroporation.

SB mediated modification of cells as described here proved to be stable in vitro and in vivo, with cells retaining transgene expression during tumor development in immunocompetent mice. The GFP+ B16F10 cells not only retained GFP expression level,but also kept a constant ratio of GFP+/GFPneg cells throughout the 15-day period of in vivo tumor development. This result suggests that no gene silencing occurs for the SB transgenic cassette, supporting in vivo utilization of this tool, as described elsewhere (Belur et al., 2003; Hausl et al., 2010).

Furthermore, we showed efficient CRISPR-mediated genome editing of *PDCD*1 gene in 293T and human PBMCs electroporated using Chicabuffers. Designing a single plasmid encoding Cas9+gRNA is simpler than constructing zinc finger nuclease (Beane et al., 2015) or TALEN (Berdien et al., 2014) based cassettes. The single plasmid approach for PBMC edition is also simpler to assemble than the recently reported Cas9+gRNA ribonucleoproteins (Schumann et al., 2015), showing that our extremely simple protocol can be used to edit cell genomes. The gRNA used for *PDCD*1 locus edition in our report targets exon 2, in contrast to exon 1 editions promoted by Schumann et al [42], showing that different gRNAs can be used to efficiently disrupt the *PDCD*1 gene sequence. The levels of gene editing obtained with our approach allow similar downstream applications in primary lymphocytes as those proposed by the above mentioned reports, but with a reduced effort to design the gene editing tool (plasmid bases CRISPR system vs TALEN or ZFN) or the electroporation reagents (plasmid vs RNA+protein). Furthermore, the protocol described for the co-electroporation of short RNAs and plasmids carrying GFP+Cas9 can be exploited for multiple loci editing in PBMCs, opening the possibility of targeting simultaneously several genes of interest.

In summary, our study describes general guidelines for the efficient electroporation of primary mammal cells and several cell lines. Furthermore, our data validates a series of flexible SB-based plasmids for the integration of transgenes and downstream selection of gene-modified cells. The combination of transposon, Chicabuffers and electroporation, as described here, represents a straightforward approach for transient gene expression and permanent gene modification of cell lines and human primary cells.

## List of abbreviations

CRISPR: clustered regularly interspaced short palindromic repeats
TALEN: Transcription activator-like effector nucleases
PBMC: peripheral blood mononuclear cells
MSC: mesenchymal stem cells
GFP: green fluorescent protein
RFP: red fluorescent protein
7-AAD: 7-amoniactinomycin D
ITR: inverted terminal repeats
SB: sleeping beauty transposase
Dpi: days post inoculation
gRNA: guide RNA
NHEJ: non-homologous end joining
FITC: Fluorescein isothiocyanate
HSC: human stem cell

## Authors’ contributions

LC, CB, BP, MC, PR and LB performed the electroporation experiments (cell lines, MSCs and PBMCs), data analysis and interpretation. CGL, CL, FRB and ZV performed electroporation and differentiation experiments in CD34+ cells, data analysis and interpretation. LC and MHB took part in the conception and design of the study, data interpretation and manuscript writing. All authors read and approved the final manuscript.

## Acknowledgments

We thank Sang Wang Han (UNIFESP – Brazil) for pT2-GFP and SB100x plasmids, Richard Morgan (NIH) for pT3-GFP plasmid and Amilcar Tanuri (UFRJ) for pRGS-CR plasmid. We also thank all the researchers that provided the cell lines used in this study.

